# Non-parametric and semi-parametric support estimation using SEquential RESampling random walks on biomolecular sequences

**DOI:** 10.1101/292078

**Authors:** Wei Wang, Jack Smith, Hussein A. Hejase, Kevin J. Liu

## Abstract

Non-parametric and semi-parametric resampling procedures are widely used to perform support estimation in computational biology and bioinformatics. Among the most widely used methods in this class is the standard bootstrap method, which consists of random sampling with replacement. While not requiring assumptions about any particular parametric model for resampling purposes, the bootstrap and related techniques assume that sites are independent and identically distributed (i.i.d.). The i.i.d. assumption can be an over-simplification for many problems in computational biology and bioinformatics. In particular, sequential dependence within biomolecular sequences is often an essential biological feature due to biochemical function, evolutionary processes such as recombination, and other factors.

To relax the simplifying i.i.d. assumption, we propose a new non-parametric/semi-parametric sequential resampling technique that generalizes “Heads-or-Tails” mirrored inputs, a simple but clever technique due to Landan and Graur. The generalized procedure takes the form of random walks along either aligned or unaligned biomolecular sequences. We refer to our new method as the SERES (or “SEquential RESampling”) method.

To demonstrate the flexibility of the new technique, we apply SERES to two different applications – one involving aligned inputs and the other involving unaligned inputs. Using simulated and empirical data, we show that SERES-based support estimation yields comparable or typically better performance compared to state-of-the-art methods for both applications.

## 1 Introduction

Resampling methods are widely used throughout computational biology and bioinformatics as a means for assessing statistical support. At a high level, resampling-based support estimation procedures consist of a methodological pipeline: resampled replicates are generated, inference/analysis is performed on each replicate, and results are then compared across replicates. Among the most widely used resampling methods are non-parametric approaches including the standard bootstrap method [Efron, 1979], which consists of random sampling with replacement. We will refer to the standard bootstrap method as the bootstrap method for brevity. Unlike parametric methods, non-parametric approaches need not assume that a particular parametric model is applicable to a problem at hand. However, the bootstrap and other widely used non-parametric approaches assume that observations are independent and identically distributed (i.i.d.).

In the context of biomolecular sequence analysis, there are a variety of biological factors that conflict with this assumption. These include evolutionary processes that cause intra-sequence dependence (e.g., recombination) and functional dependence among biomolecular sequence elements and motifs. Felsenstein presciently noted these limitations when he proposed the application of the bootstrap to phylogenetic inference: “A more serious difficulty is lack of independence of the evolutionary processes in different characters. … For the purposes of this paper, we will ignore these correlations and assume that they cause no problems; in practice, they pose the most serious challenge to the use of bootstrap methods.” (reproduced from p. 785 of [Felsenstein, 1985]).

To relax the simplifying assumption of i.i.d. observations, Landan and Graur [2007] introduced the Heads-or-Tails (HoT) technique for the specific problem of multiple sequence alignment (MSA) support estimation. The idea behind HoT is simple but quite powerful: inference/analysis should be repeatable whether an MSA is read either from left-to-right or from right-to-left – i.e., in either heads or tails direction, respectively. While HoT resampling preserves intra-sequence dependence, it is limited to two replicates, which is far fewer than typically needed for reasonable support estimation; often, hundreds of resampled replicates or more are used in practice. Subsequently developed support estimation procedures increased the number of possible replicates by augmenting HoT with bootstrapping, parametric resampling, and domain-specific techniques (e.g., progressive MSA estimation) [Landan and Graur, 2008, Penn et al., 2010, Sela et al., 2015]. The combined procedures were shown to yield comparable or improved support estimates relative to the original HoT procedure [Sela et al., 2015] as well as other state-of-the-art parametric and domain-specific methods [Kim and Ma, 2011, Notredame et al., 2000], at the cost of some of the generalizability inherent to non-parametric approaches. In this study, we revisit the central question that HoT partially addressed: how can we resample many non-parametric replicates that account for dependence within a sequence of observations, and how can such techniques be used to derive improved support estimates for biomolecular sequence analysis?

## 2 Methods

In our view, a more general statement of HoT’s main insight is the following, which we refer to as the “neighbor preservation property”: a neighboring observation is still a neighbor, whether reading an observation sequence from the left or the right. In other words, the key property needed for non-parametric resampling is preservation of neighboring bases within the original sequences, where any pair of bases that appear as neighbors in a resampled sequence must also be neighbors in the corresponding original sequence. To obtain many resampled replicates that account for intra-sequence dependence while retaining the neighbor preservation property, we propose a random walk procedure which generalizes a combination of the bootstrap method and the HoT method. We refer to the new resampling procedure as SERES (“SEquential RESampling”). Note that the neighbor preservation property is necessary but not sufficient for statistical support estimation. Other important properties include computational efficiency of the resampling procedure and unbiased sampling of observations within the original observation sequence.

SERES walks can be performed on both aligned and unaligned sequence inputs. We discuss the case of aligned inputs first, since it is simpler than the case of unaligned inputs.

### 2.1 SERES walks on aligned sequences

Detailed pseudocode for a non-parametric SERES walk on a fixed MSA is shown in the Appendix’s Supplementary Methods section: Algorithm 1.

The random walk is performed on the sequence of aligned characters (i.e., MSA sites). The starting point for the walk is chosen uniformly at random from the alignment sites, and the starting direction is also chosen uniformly at random. The random walk then proceeds in the chosen direction with non-deterministic reversals, or direction changes, that occur with probability γ; furthermore, reversals occur with certainty at the start and end of the fixed MSA. Aligned characters are sampled during each step of the walk. The random walk ends once the number of sampled characters is equal to the fixed MSA length.

The long-term behavior of an infinitely long SERES random walk can be described by a second-order Markov chain. Certain special cases (e.g., γ = 0.5) can be described using a first-order Markov chain.

In theory, a finite-length SERES random walk can exhibit biased sampling of sites since reversal occurs with certainty at the start and end of the observation sequence, whereas reversal occurs with probability γ elsewhere. However, for practical choices of walk length and reversal probability γ, sampling bias is expected to be minimal.

### 2.2 SERES walks on unaligned sequences

Detailed pseudocode for SERES resampling of unaligned sequences is shown in the Appendix’s Supplementary Methods section: see Algorithm 2. Figure 1 provides an illustrated example.

The procedure begins with estimating a set of anchors – sequence regions that exhibit high sequence similarity – which enable resampling synchronization across unaligned sequences. A conservative approach for identifying anchors would be to use highly similar regions that appear in the strict consensus of multiple MSA estimation methods. In practice, we found that highly similar regions within a single guide MSA produced reasonable anchors. We used the average normalized Hamming distance (ANHD) as our similarity measure, where indels are treated as mismatches.

Unaligned sequence indices corresponding to the start and end of each anchor serve as “barriers” in much the same sense as in parallel computing: asynchronous sequence reads occur between barrier pairs along a current direction (left or right), and a random walk is conducted on barrier space in a manner similar to a SERES walk on a sequence of aligned characters. The set of barriers also includes trivial barriers at the start and end of the unaligned sequences. The random walk concludes once the unaligned sequences in the resampled replicate have sufficient length; our criterion requires that the longest resampled sequence has minimum length that is a multiple maxReplicateLengthFactor of the longest input sequence length.

Technically, the anchors used in our study make use of parametric MSA estimation and the rest of the SERES walk is non-parametric. The overall procedure is therefore semi-parametric (although see Conclusions for an alternative).

### 2.3 Performance study overview

Our study evaluated the performance of SERES-based support estimation in the context of two applications – one utilizing fixed alignments as input and the other utilizing unaligned sequences as input. The aligned input application is posterior decoding of phylo-HMMs for the task of analyzing local genealogical variation due to recombination. The unaligned input application is MSA support estimation. Of course, there are many other applications for non-parametric support estimation – too many to investigate in one study. We focus on the above two applications since they cover the two different classes of SERES inputs. Furthermore, the two applications are considered to be classical problems in computational biology and bioinformatics and their outputs are useful for studying a range of topics (e.g., phylogenetics and phylogenomics, proteomics, comparative genomics, etc.).

### 2.4 Aligned input application: posterior decoding of phylo-HMMs

#### Computational methods

The coalescent-with-recombination (CwR) model [Hudson, 1983] is a classical population genetic model involving recombination. However, phylogenetic inference under the multi-species CwR model is computationally prohibitive, and alternatives such as the sequentially Markovian coalescent (SMC) model [McVean and Cardin, 2005] are used as an approximation to the full CwR model. First-order hidden Markov models (HMMs) are a widely used choice for tractable SMC-based inference. Phylo-HMMs are the class of HMMs with hidden states that correspond to phylogenies. Markovian dependence between phylo-HMM states are meant to capture intra-sequence dependence among local phylogenies, which can be caused by recombination and other evolutionary processes. There are a variety of HMM-based methods for local genealogical inference, depending on modeling assumptions [Husmeier and Wright, 2001, Westesson and Holmes, 2009, Mailund et al., 2011, Liu et al., 2014]. We focus on recHMM [Westesson and Holmes, 2009] as an exemplar method in this class. We ran recHMM with default settings. Consistent with the study of Westesson and Holmes [2009], we used the posterior decoding algorithm to perform statistical inference of local phylogenies [Rabiner, 1989]. The posterior decoding algorithm addresses the following problem. Let *G* be the set of all possible unrooted tree topologies on *n* taxa. The input consists of a multiple sequence alignment *A* on *n* sequences – one for each of *n* taxa – with length *k* (i.e., *k* sites in *A*). *A* is assumed to contain recombinant sequences, and historical recombination can cause local genealogies to vary across the sites in *A* [Hein et al., 2004]. The output consists of the following: for each aligned site *a*_*i*_ where 1 ≤ *i* ≤ *k*, we seek the conditional probability that the HMM is in a hidden state corresponding to a particular gene tree *g* ∈ *G* conditional on all sites in *A* and the fitted HMM model. For a particular HMM instance, the posterior decoding effectively estimates which gene tree is the most likely evolutionary history that explains the observed character at a given site conditional on the sequence of all observed sites in *A*. Analogous to the distinction between filtering and smoothing, the posterior decoding weighs any particular inference at a given site against the total evidence across all sites.

**Fig. 1.**
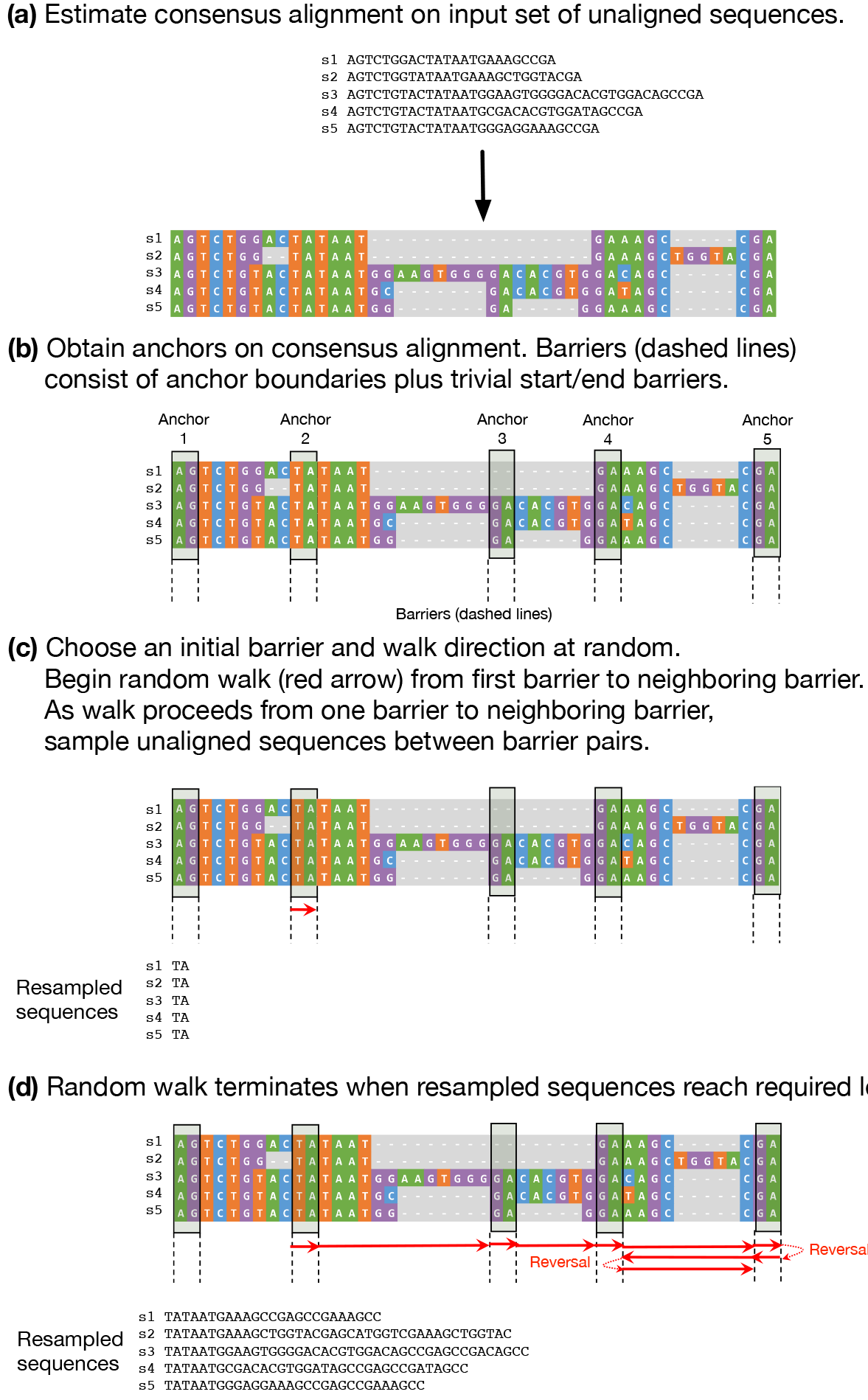
Illustrated example of SERES resampling random walk on unaligned sequences. Detailed pseudocode is provided in the Appendix’s Supplementary Methods section (Algorithm 2 in Appendix), (a) The resampling procedure begins with the estimation of a consensus alignment on the input set of unaligned sequences, (b) A set of conservative anchors is then obtained using the consensus alignment, and anchor boundaries define a set of barriers (including two trivial barriers – one at the start of the sequences and one at the end of the sequences), (c) The SERES random walk is conducted on the set of barriers. The walk begins at a random barrier and proceeds in a random direction to the neighboring barrier. The walk reverses with certainty when the trivial start/end barriers are encountered; furthermore, the walk direction can randomly reverse with probability γ. As the walk proceeds from barrier to barrier, unaligned sequences are sampled between neighboring barrier pairs, (d) The resampling procedure terminates when the resampled sequences meet a specified sequence length threshold.

We used SERES resampling and recHMM as part of a support estimation pipeline. First, we ran SERES resampling on the input alignment *A* (Algorithm 1 in Appendix). The SERES resampling procedure used reversal probability γ = 0.005. Thirty SERES replicates were generated per simulated dataset. Next, we ran recHMM using default settings on each SERES replicate, and the posterior decoding algorithm was used to infer posterior decoding probabilities for each site. For each site, inferred posterior decoding probability distributions were aggregated across all SERES replicates in which the site appeared (with per-replicate multiplicity based on the number of times that the site was sampled within the replicate). The aggregated distribution was then normalized to obtain a valid probability distribution. Finally, we used the peak of the distribution and its corresponding HMM state and local topology as our inference. To facilitate comparison, the same peak-based inference procedure was used with recHMM-based posterior decoding of the original input alignment.

#### Simulated datasets

Gene trees were simulated under the CwR model using ms [Hudson, 2002]. Each CwR simulation sampled either 4 or 6 alleles with scaled recombination rate *ρ* ∈ {0.5,1.0, 2.0} and total sequence length per replicate of 1 kb. For each gene tree, finite-length sequence evolution was simulated under the Jukes-Cantor model of nucleotide substitution [Jukes and Cantor, 1969] using Seq-Gen [Rambaut and Grassly, 1997]. We used a mutation rate *θ* ∈ {0.5,1.0, 2.0}. A model condition consisted of fixed values for all model parameters, and simulation procedures were repeated so that 30 replicate datasets were generated per model condition. We assessed topological accuracy of inferred gene trees relative to ground truth using the Robinson-Foulds measure [Robinson and Foulds, 1981], which is the proportion of bipartitions that occur in an inferred gene tree but not the true gene tree or vice versa.

### 2.5 Unaligned input application: MSA support estimation

#### Computational methods

We examined the problem of evaluating support in the context of MSA estimation. The problem input consists of an estimated MSA *A* which has a corresponding set of unaligned sequences *S*. The problem output consists of support estimates for each nucleotide-nucleotide homology in *A*, where each support estimate is on the unit interval. Note that this computational problem is distinct from the full MSA estimation problem.

There are a variety of existing methods for MSA support estimation. The creators of HoT and their collaborators subsequently developed alignment-specific parametric resampling techniques [Landan and Graur, 2008] and then combined the two to obtain two new semi-parametric approaches: GUIDANCE [Penn et al., 2010] (which we will refer to as GUIDANCE1) and GUIDANCE2 [Sela et al., 2015]. Other parametric MSA support estimation methods include PSAR [Kim and Ma, 2011] and T-Coffee [Notredame et al., 2000].

We focus on GUIDANCE1 and GUIDANCE2, which have been demonstrated to have comparable or better performance relative to other state-of-the-art methods [Sela et al., 2015]. We used MAFFT for re-estimation on resampled replicates, since it has been shown to be among the most accurate progressive MSA methods to date [Katoh and Standley, 2013, Liu et al., 2012].

We then used SERES to perform resampling in place of the standard bootstrap that is used in the first step of GUIDANCE1/GUIDANCE2. Re-estimation was performed on 100 SERES replicates – each consisting of a set of unaligned sequences – using MAFFT with default settings, which corresponds to the FFT-NS-2 algorithm for progressive alignment. The SERES resampling procedure used a reversal probability γ = 0.5, which is equivalent to selecting a direction uniformly at random (UAR) at each step of the random walk; each SERES replicate utilized a total of 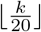 anchors with anchor size of 5 bp and a minimum distance between neighboring anchors of 25 bp, where *k* is the length of the input alignment *A*. All downstream steps of GUIDANCE1/GUIDANCE2 were then performed using the re-estimated alignments as input.

#### Simulated datasets

Model trees and sequences were simulated using INDELible [Fletcher and Yang, 2009]. First, non-ultrametric model trees with either 10 or 50 taxa were sampled using the following procedure: model trees were generated under a birth-death process [Yang and Rannala, 1997], branch lengths were chosen UAR from the interval (0,1), and the model tree height was re-scaled from its original height *h*_0_ to a desired height *h* by multiplying all branch lengths by the factor *h*/*h*_0_. Next, sequences were evolved down each model tree under the General Time-Reversible (GTR) model of substitution [Rodriguez et al., 1990] and the indel model of Fletcher and Yang [2009], where the root sequence had length of 1 kb. We used the substitution rates and base frequencies from the study of Liu et al. [2012], which were based upon empirical analysis of the nematode Tree of Life. Sequence insertions/deletions occurred at rate *r*_*i*_, and we used the medium gap length distribution from the study of Liu et al. [2012]. The model parameter values used for simulation are shown in the Appendix (Supplementary Table S1), and each combination of model parameter values constitutes a model condition. Model conditions are enumerated in order of generally increasing sequence divergence, as reflected by average pairwise ANHD. For each model condition, the simulation procedure was repeated to generate twenty replicate datasets. Summary statistics for simulated datasets are shown in Supplementary Table S1.

We evaluated performance based upon receiver operating character (ROC) curves, precision-recall curves (PR), and area under ROC and PR curves (ROC-AUC and PR-AUC, respectively). Consistent with other studies of MSA support estimation techniques [Penn et al., 2010, Sela et al., 2015], the MSA support estimation problem in our study entails annotation of nucleotide-nucleotide homologies in the estimated alignment; thus, homologies that appear in the true alignment but not the estimated alignment are not considered. For this reason, the confusion matrix quantities used for ROC and PR calculations are defined as follows. True positives (TP) are the set of nucleotide-nucleotide homologies that appear in the true alignment and the estimated alignment with support value greater than or equal to a given threshold, false positives (FP) are the set of nucleotide-nucleotide homologies that appear in the estimated alignment with support value greater than or equal to a given threshold but do not appear in the true alignment, false negatives (FN) are the set of nucleotide-nucleotide homologies that appear in the true alignment but appear in the estimated alignment with support value below a given threshold, and true negatives (TN) are the set of nucleotide-nucleotide homologies that do not appear in the true alignment and appear in the estimated alignment with support value below a given threshold. The ROC curve plots the true positive rate (|TP|/(|TP| + |FN|)) versus the false positive rate (|FP|/(|FP| + |TN|)). The PR curve plots the true positive rate versus precision (|TP|/(|TP| + |FP|)). Varying the support threshold yields different points along these curves. Custom scripts were used to perform confusion matrix calculations. ROC curve, PR curve, and AUROC calculations were performed using the scikit-learn Python library [Pedregosa et al., 2011].

#### Empirical datasets

We downloaded empirical benchmarks from the Comparative RNA Web (CRW) Site database, which can be found at www.rna.icmb.utexas.edu [Cannone et al., 2002]. In brief, the CRW database includes riboso-mal RNA sequence datasets than span a range of dataset sizes and evolutionary divergence. We focused on datasets where high-quality reference alignments are available; the reference alignments were produced using intensive manual curation and analysis of heterogeneous data, including secondary structure information. We selected primary 16S rRNA, primary 23S rRNA, primary intron, and seed alignments with at most 250 sequences. Aligned sequences with 99% or more missing data and/or indels were omitted from analysis. Summary statistics for the empirical benchmarks are shown in the Appendix (Supplementary Table S2).

### 2.6 Software and data availability

Open-source software and open data can be found at https://gitlab.msu.edu/liulab/SERES-Scripts-Data.

## 3 Results

### 3.1 Aligned input application: posterior decoding of phylo-HMMs

#### Simulation study

Relative to standalone posterior decoding, the SERES-based method yielded statistically significant improvements in topological accuracy on all model conditions with one exception (Figure 2). The single exception occurred on the smallest model condition with the lowest recombination rate and mutation rate in our study (both 0.5), where a small but statistically insignificant improvement was seen.

On the four-taxon model conditions with the lowest recombination rate (*ρ* = 0.5), the SERES-based method’s advantage in topological accuracy grew as the mutation rate increased from 0.5 to 2. On all other four-taxon model conditions, the SERES-based method’s average improvement in topological accuracy over standalone posterior decoding was mostly unchanged as the recombination rate and mutation rate varied. A similar outcome was observed on the six-taxon model conditions, where the difference in average topological error between the two methods was generally similar across a range of recombination rates and mutation rates. Overall, topological error was generally greater on the six-taxon model conditions compared to otherwise equivalent four-taxon model conditions. The quantitative improvement seen on the six-taxon model conditions was generally less than on the four-taxon model conditions as well. The largest improvements were seen on the four-taxon model conditions with recombination rate *ρ* = 2, which were as large as 18.7%.

### 3.2 Unaligned input application: MSA support estimation

#### Simulation study

For all model conditions, SERES-based resampling and reestimation yielded improved MSA support estimates compared to GUIDANCE1 and GUIDANCE2, two state-of-the-art methods, where performance was measured by PR-AUC or ROC-AUC (Table 1). In all cases, PR-AUC or ROC-AUC improvements were statistically significant (corrected pairwise t-test or DeLong et al. [1988] test, respectively; *n* = 20 and *α* = 0.5). The observed performance improvement was robust to several experimental factors: dataset size, increasing sequence divergence due to increasing numbers of substitutions, insertions, and deletions, and the choice of alignment-specific parametric support estimation techniques (i.e., the parametric approaches used by either GUIDANCE1 or GUIDANCE2) that were used in combination with SERES-based support estimation.

Compared to dataset size, sequence divergence had a relatively greater quantitative impact on each method’s performance. For each dataset size (10 or 50 taxa), PR-AUC differed by at most 3% on the least divergent model condition. The SERES-based method’s performance advantage grew as sequence divergence increased – to as much as 28% – and the largest performance advantages were seen on the most divergent datasets in our study. The most divergent datasets were also the most challenging. For each method, PR-AUC generally degraded as sequence divergence increased; however, the SERES-based method’s PR-AUC degraded more slowly compared to the non-SERES-based method. Consistent with the study of Sela et al. [2015], GUIDANCE2 consistently outperformed GUIDANCE1 on each model conditions and using either AUC measure. The performance improvement of SERES+GUIDANCE1 over GUIDANCE1 was generally greater than that seen when comparing SERES+GUIDANCE2 and GUIDANCE2; furthermore, the PR-AUC-based corrected q-values were more significant for the former compared to the latter in all cases except for the 10.D model condition, where the corrected q-values were comparable. Finally, while the SERES-based method consistently yielded performance improvements over the corresponding non-SERES-based method regardless of the choice of performance measure (either PR-AUC or ROC-AUC), the PR-AUC difference was generally larger than the ROC-AUC difference, especially on more divergent model conditions.

**Fig. 2.**
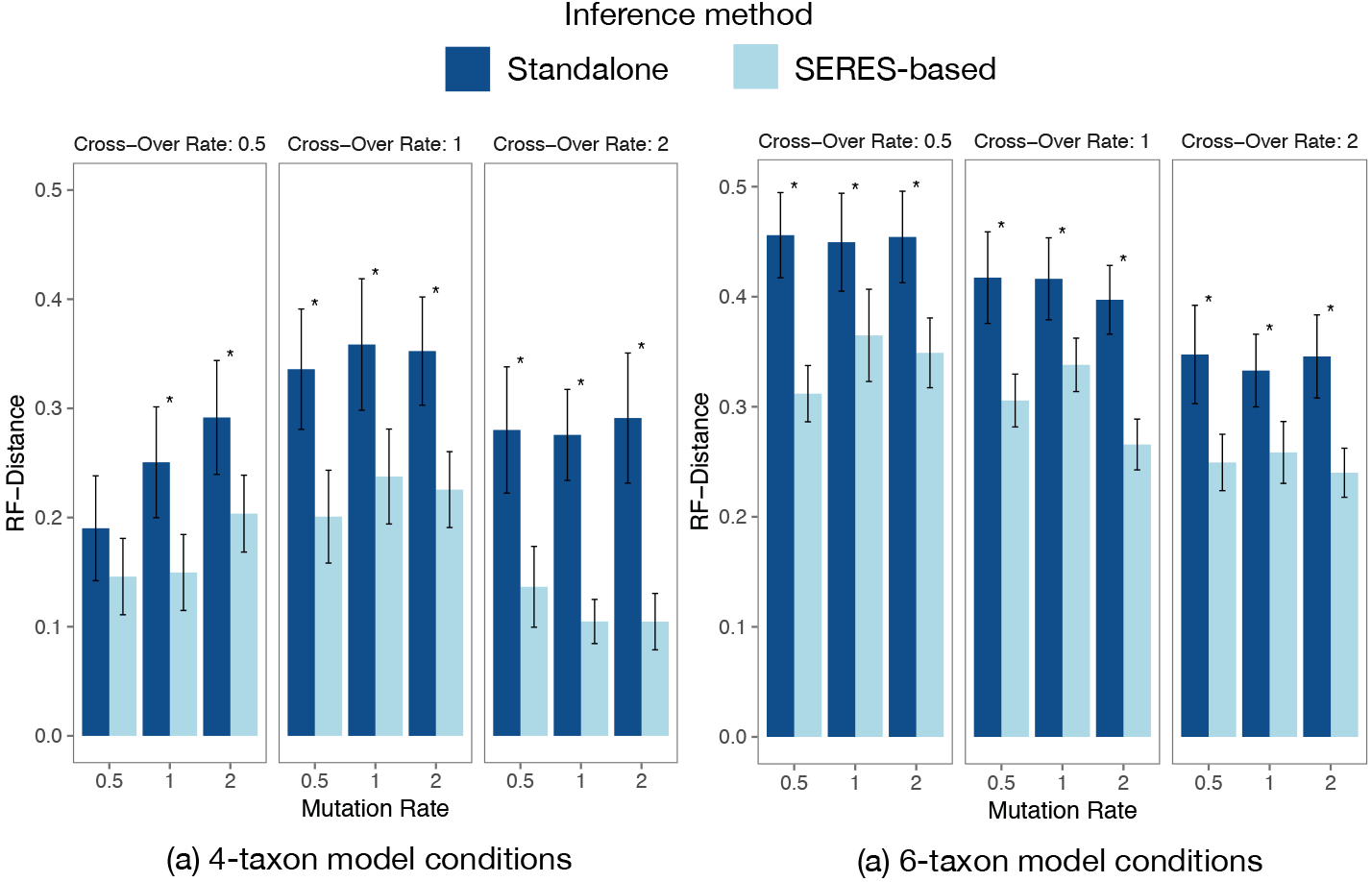
Aligned input application: simulation study results. Results are shown for model conditions with either 4 or 6 taxa, recombination rate between 0.5 and 2.0, and mutation rate between 0.5 and 2.0. The topological error of local gene trees inferred using SERES-based posterior decoding (light blue) is compared to standalone posterior decoding (dark blue). Topological error is measured using Robinson-Foulds distance [Robinson and Foulds, 1981] between the inferred and true gene trees, where we report each method’s average topological error across all sites and replicates in a model condition. Average and standard error bars are shown (*n* = 30). To test whether SERES-based inference yielded a significant improvement in topological error over standalone posterior decoding, we performed a one-tailed pairwise t-test with multiple test correction using the Benjamini-Hochberg method [Benjamini and Hochberg, 1995] (*α* = 0.05 and *n* = 30); asterisks denote statistical significance based upon the corrected test.

#### Empirical study

Relative to GUIDANCE1 or GUIDANCE2, SERES-based support estimates consistently returned higher AUC on all datasets – primary, seed, and intronic – with a single exception: the comparison of SERES+GUIDANCE2 and GUIDANCE2 on the intronic IGIC2 dataset, where the PR-AUC and ROC-AUC differences were 1.17% and 2.12%, respectively. For each pairwise comparison of methods (i.e., SERES+GUIDANCE1 vs. GUIDANCE1 or SERES+GUIDANCE2 vs. GUIDANCE2), the SERES-based method returned relatively larger PR-AUC improvements on datasets with greater sequence divergence, as measured by ANHD and gappiness. In particular, PR-AUC improvements were less than 1% on seed and primary non-intronic datasets. Intronic datasets yielded PR-AUC improvements of as much as 13.87%. Observed AUC improvements of SERES+GUIDANCE1 over GUIDANCE1 were relatively greater than those seen for SERES+GUIDANCE2 in comparison to GUIDANCE2. Finally, GUIDANCE2 consistently returned higher AUC relative to GUIDANCE1, regardless of whether PR or ROC curves were the basis for AUC comparison.

## 4 Discussion

Re-estimation using SERES resampling resulted in comparable or typically improved support estimates for the applications in our study. We believe that this performance advantage is due to the ability to generate many distinct replicates while enforcing the neighbor preservation principle. The latter is critical for retaining sequence dependence which is inherent to both applications in our study.

### Aligned input application: posterior decoding of phylo-HMMs

For all model conditions, recHMM re-estimation using SERES resampling returned improved average topological error compared to standalone recHMM analysis. The improvement was statistically significant for all model conditions except the model condition with the smallest number of taxa and lowest mutation rate and recombination rate in our study. Standalone recHMM analysis was most accurate on the latter model condition relative to more divergent and larger model conditions. Furthermore, we found that the methods in our study had comparably low inference error on this model condition. These findings suggest that the reduced input size and sequence divergence of this model condition may have posed less of a challenge for the purposes of inference.

**Table 1.**
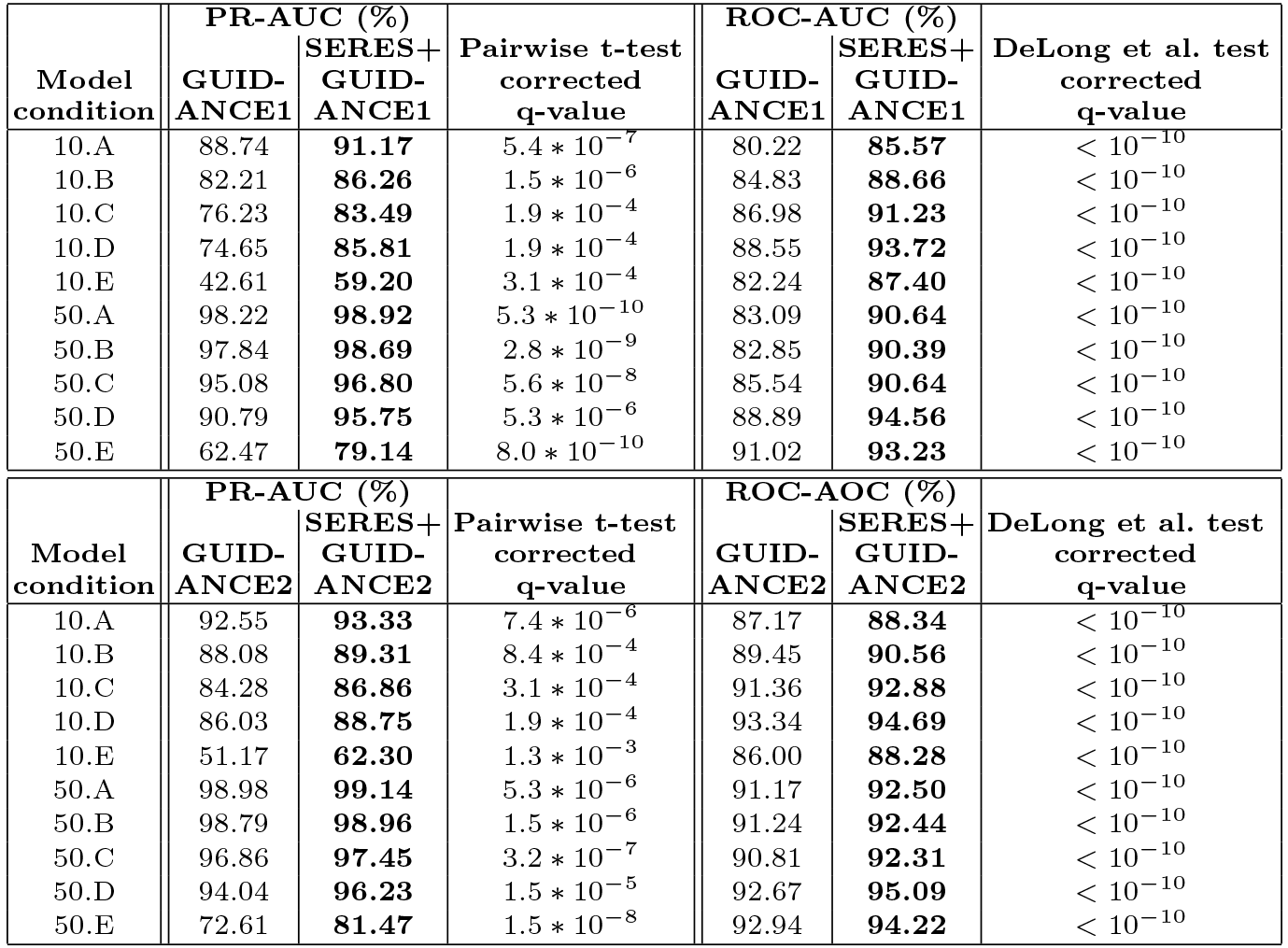
Unaligned input application: simulation study results. Results are shown for five 10-taxon model conditions (named 10.A through 10.E in order of generally increasing sequence divergence) and five 50-taxon model conditions (similarly named 50.A through 50.E). We evaluated the performance of two state-of-the-art methods for MSA support estimation – GUIDANCE1 [Penn et al., 2010] and GUIDANCE2 [Sela et al., 2015] – versus re-estimation on SERES and parametrically resampled replicates (using parametric techniques from either GUIDANCE1 or GUIDANCE2). (See Methods section for details.) We calculated each method’s precision-recall (PR) and receiver operating characteristic (ROC) curves. Performance is evaluated based upon aggregate area under curve (AUC) across all replicates for a model condition (*n* = 20). The top rows show AUC comparisons of GUIDANCE1 (“GUIDANCE1”) vs. SERES combined with parametric techniques from GUIDANCE1 (“SERES+GUIDANCE1”), and the bottom rows show AUC comparisons of GUIDANCE2 (“GUIDANCE2”) vs. SERES combined with parametric techniques from GUIDANCE2 (“SERES+GUIDANCE2”); for each model condition and pairwise comparison, the best AUC is shown in bold. Statistical significance of PR-AUC or AUC-ROC differences was assessed using a one-tailed pairwise t-test or DeLong et al. [1988] test, respectively, and multiple test correction was performed using Benjamini and Hochberg [1995]’s method. Corrected q-values are reported (*n* = 20) and all were significant (*α* = 0.05).

**Table 2.**
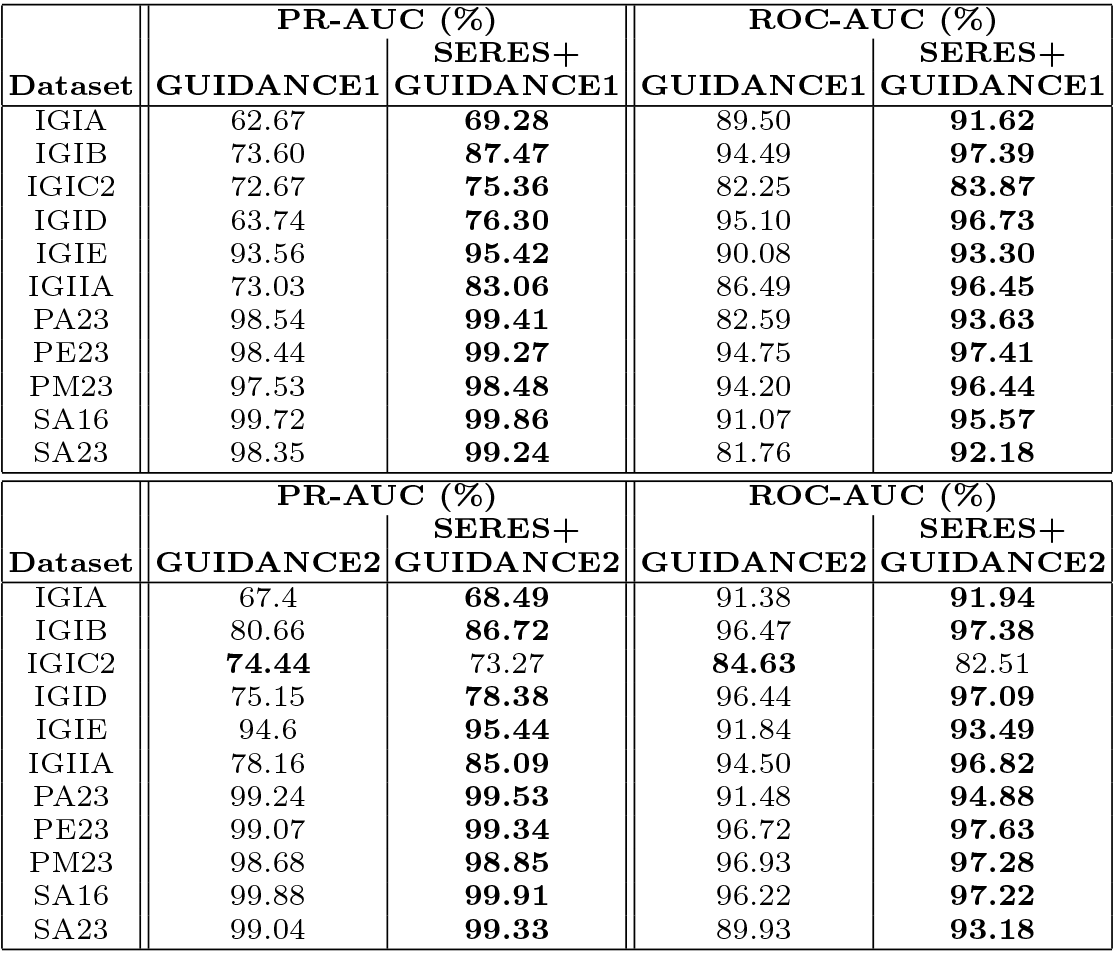
Unaligned input application: empirical study results. The empirical study made use of benchmark RNA datasets and curated reference alignments from the CRW database [Cannone et al., 2002]. Results are shown for intronic (“IG” prefix) and non-intronic datasets (“P” prefix and “S” prefix, following “primary” and “seed” nomenclature from the CRW database). For each dataset, we report each method’s PR-AUC and ROC-AUC. For each dataset and pairwise method comparison, the best AUC is shown in bold. Methods, performance measures, table layout, and table description are otherwise identical to Table 1.

### Unaligned input application: MSA support estimation

On all model conditions, SERES+GUIDANCE1 support estimation resulted in significant improvements in AUC-PR and AUC-ROC compared to GUIDANCE1. A similar outcome was observed when comparing SERES+GUIDANCE2 and GUIDANCE2. The main difference in each comparison is the resampling technique – either SERES or standard bootstrap. Our findings clearly demonstrate the performance advantage of the former over the latter. SERES accounts for intrasequence dependence due to insertion and deletion processes, while the bootstrap method assumes that sites are independent and identically distributed. Regarding comparisons involving GUIDANCE2 versus GUIDANCE1, a contributing factor may have been the greater AUC of GUIDANCE2 over GUIDANCE1. We used SERES to perform semi-parametric support estimation in conjunction with the parametric support techniques of GUIDANCE1 or GUIDANCE2. The latter method’s relatively greater AUC may be more challenging to improve upon.

The performance comparisons on empirical benchmarks were consistent with the simulation study. In terms of ANHD and gappiness, the non-intronic datasets in our empirical study were more like the low divergence model conditions in our simulation study, and the intronic datasets were more like the higher divergence model conditions. Across all empirical datasets, SERES-based support estimation consistently yielded comparable or better AUC versus GUIDANCE1 or GUIDANCE2 alone. The SERES-based method’s AUC advantage generally increased as datasets became more divergent and challenging to align – particularly when comparing performance on non-intronic versus intronic datasets. We found that the support estimation methods returned comparable AUC (within a few percentage points) on datasets with 1-2 dozen sequences and low sequence divergence relative to other datasets. In particular, IGIC2 was the only dataset where SERES+GUIDANCE2 did not return an improved AUC relative to GUIDANCE2. IGIC2 was the second-smallest dataset – about an order of magnitude smaller than all other datasets except the IGID dataset – and IGIC2 also had the second-lowest ANHD and lowest gappiness among intronic datasets. IGID was the smallest dataset, but had higher ANHD and gappiness compared to the IGIC2 dataset. Compared to the other empirical datasets, SERES+GUIDANCE2 returned a small AUC improvement over GUIDANCE2 on the IGID dataset – at most 3.2%.

On simulated and empirical datasets, greater sequence divergence generally resulted in increased inference error for all methods. However, the SERES-based method’s performance tended to degrade more slowly than the corresponding non-SERES-based method as sequence divergence increased, and the greatest performance advantage was seen on the most divergent model conditions and empirical datasets.

Finally, we note that non-parametric/semi-parametric resampling techniques are orthogonal to parametric alternatives. Consistent with previous studies [Penn et al., 2010, Sela et al., 2015], we found that combining two different classes of methods yielded better performance than either by itself.

## 5 Conclusion

This study introduced SERES, which consists of new non-parametric and semi-parametric techniques for resampling biomolecular sequence data. Using simulated and empirical data, we explored the use of SERES resampling for support estimation in two classical problems in computational biology and bioinformatics – one involving aligned sequences and the other involving unaligned sequences. We found that SERES-based support estimation yields comparable or typically better performance compared to state-of-the-art approaches.

We conclude with possible directions for future work. The SERES algorithm in our study made use of a semi-parametric resampling procedure on unaligned inputs, since anchors were constructed using progressive multiple sequence alignment. While this approach worked well in our experiments, non-parametric alternatives could be substituted (e.g., unsupervised *k*-mer clustering using alignment-free distances [Daskalakis and Roch, 2010]) to obtain a purely non-parametric resampling procedure. Second, the unaligned input application focused on nucleotide-nucleotide homologies to enable direct comparison against existing MSA support estimation procedures (i.e., GUIDANCE1 and GUIDANCE2). The SERES framework can be extended in a straightforward manner to estimate support for nucleotide-indel pairs. Third, SERES resampling can be used to perform full MSA inference. A simple approach would be to analyze homologies that appeared in re-estimated inferences across resampled replicates, without regard to any input alignment. Finally, we envision many other SERES applications. Examples in computational biology and bioinformatics include protein structure prediction, detecting genomic patterns of natural selection, and read mapping and assembly. Non-parametric resampling for support estimation is widely used throughout science and engineering, and SERES resampling may similarly prove useful in research areas outside of computational biology and bioinformatics.

## Acknowledgments

This work has been supported in part by the National Science Foundation (grant nos. CCF-1565719, CCF-1714417, and DEB-1737898) and MSU faculty startup funds (to KJL). Computational experiments were performed using the High Performance Computing Center (HPCC) at Michigan State University (MSU).

